# Solart: A Structure-Based Method To Predict Protein Solubility And Aggregation

**DOI:** 10.1101/600734

**Authors:** Q. Hou, J. M. Kwasigroch, M. Rooman, F. Pucci

## Abstract

**Motivation:** The solubility of a protein is often decisive for its proper functioning. Lack of solubility is a major bottleneck in high-throughput structural genomic studies and in high-concentration protein production, and the formation of protein aggregates causes a wide variety of diseases. Since solubility measurements are time-consuming and expensive, there is a strong need for solubility prediction tools.

**Results:** We have recently introduced solubility-dependent distance potentials that are able to unravel the role of residue-residue interactions in promoting or decreasing protein solubility. Here, we extended their construction by defining solubility-dependent potentials based on backbone torsion angles and solvent accessibility, and integrated them, together with other structure- and sequence-based features, into a random forest model trained on a set of *E. coli* proteins with experimental structures and solubility values. We thus obtained the SOLart protein solubility predictor, whose most informative features turned out to be folding free energy differences computed from our solubility-dependent statistical potentials. SOLart performances are very good, with a Pearson correlation coefficient between experimental and predicted solubility values of 0.7 both in the training dataset and on an independent set of *S. Cerevisiae* proteins. On test sets of modeled structures, only a limited drop in performance is observed. SOLart can thus be used with both high-resolution and low-resolution structures, and clearly outperforms state-of-art solubility predictors. It is available through a user-friendly webserver, which is easy to use by non-expert scientists.

**Availability:** The SOLart webserver is freely available at babylone.ulb.ac.be/SOLART/

## 1. Introduction

Solubility and aggregation are crucial properties of proteins, which can either ensure or prevent their correct functioning (Trevino *et al.*, 2008). Obtaining a thorough understanding of these matters is becoming increasingly important, since protein solubilization is required for improving a wide range of biotechnological and biopharmaceutical processes, especially when high protein concentrations are demanded. Just to mention some of them, protein solubility is frequently a serious bottleneck for the successful development of antibody therapeutics, which often suffer from aggregation at the conditions in which they are stored (Perchiacca and Tessier, 2012; Roberts, 2014), as well as in genome-wide structural analyses where about 80% of the total number of non-membrane proteins have been estimated to have insolubility-related problems (Golovanov *et al.*, 2004).

In the context of recombinant protein production, the formation of insoluble inclusion bodies, which are thought to contain clusters of different conformational states corresponding to folded, misfolded and partially folded structures, frequently makes the procedure to get bioactive proteins very challenging, involving first the solubilization of the inclusion bodies followed by the native refolding of the proteins (Martínez-Alonso *et al.*, 2009; Singh *et al.*, 2015; Baneyx and Mujacic, 2004; Singh and Panda, 2005; Vallejo and Rinas, 2004).

Scarse solubility properties are also directly related to pathological conditions such as the neurodegenerative Alzheimer and Parkinson diseases, whose hallmark is the progressive accumulation of insoluble deposits, *i.e β*-amyloid and *α*-synuclein aggregates, respectively, that become toxic and interfere with the normal cell functioning (Chiti and Dobson, 2006; Bucciantini *et al.*, 2002; Irvine *et al.*, 2008; Ross and Poirier, 2004).

Reaching a full understanding of protein solubility mechanisms is particularly challenging, since solubility is a complex physicochemical property determined not only by various intrinsic factors such as residue-residue interactions, protein flexibility, amino acid composition and hydrophobicity, but also by various extrinsic variables such as the pH, the environmental temperature, the ionic strength and the protein concentration.

During the past decades, many efforts have been devoted to investigate the mechanisms and the factors that influence protein solubility (Trainor *et al.*, 2017). It has been reported that smaller proteins tend to have a higher solubility when overexpressed in *E. Coli* than longer proteins (Wilkinson and Harrison, 1991). The amino acid composition also influences protein solubility. For example, Asp, Glu and Ser contribute more favorably to solubility than other hydrophilic amino acids (Niwa *et al.*, 2009; Chan *et al.*, 2013); the values of the Lys/Arg and Glu/Asp ratios correlate with solubility (Warwicker *et al.*, 2013; Chan *et al.*, 2013); and aromatic-poor proteins tend to be more soluble than those enriched in aromatics (Niwa *et al.*, 2009; Hebditch *et al.*, 2017).

Furthermore, protein-protein and protein-solvent interactions have been shown to play key roles in the solubility properties. In particular, solvent exposed residues have some characteristics that are well correlated with solubility: insoluble proteins tend to have larger surface patches carrying a net positive charge than soluble proteins (Chan *et al.*, 2013), which are characterized instead by a more negatively charged surface (Kramer *et al.*, 2012).

In a recent work (Hou *et al.*, 2018), we showed that among all residue-residue interactions, the Lys-containing salt bridges and the aliphatic interactions contribute more strongly than others to promote solubility, whereas interactions involving delocalized *π*-electrons favor aggregation (*e.g.* aromatic, His-*π*, cation-*π*, amino-*π* and anion-*π* interactions). These different findings demonstrate the important potentiality of structural information in the understanding of the biophysical mechanisms underlying solubility data.

Several computational methods, mostly based on machine learning techniques, have recently been developed to predict protein solubility (Smialowski *et al.*, 2006, 2012; Idicula-Thomas *et al.*, 2005; Magnan *et al.*, 2009; Agostini *et al.*, 2014; Hebditch *et al.*, 2017; Khurana *et al.*, 2018; Hirose and Noguchi, 2013; Sormanni *et al.*, 2015). The large majority of the features that they use are extracted from the amino acid sequences, such as the sequence length, the amino acid composition, the absolute charge, the isoelectric point, the aliphatic index and the average hydropathy. Some other features are associated to structural properties, such as *β*-stand propensities or fractions of exposed and buried residues. However, these features are not assigned from the structure, but rather predicted from the sequence.

Structure-based techniques to predict solubility make use of extensive molecular dynamics simulations to evaluate the free energy difference between solution and aggregation phases (Tjong and Zhou, 2008). However these methods are computationally expensive and cannot be certainly applied to large-scale investigations of the protein solubility.

In summary, the current prediction methods tend to overlook structural data and require only the sequence as input. Clearly, considering features derived from experimental 3-dimensional (3D) structures adds valuable information, which should in principle boost the methods’ performances. However, requiring 3D structures decreases the applicability of the predictor, as they are not always available. But this drawback is loosing importance, since homology modeling tools provide always better structural models that can safely be used by some predictors. Hence, progress is definitely expected in the solubility prediction field from the utilization of 3D structures.

In this paper, we fully exploited protein structure data through the use of statistical potentials, which have largely proven to be successful in many studies ranging from structure prediction to mutant analyses (see *e.g.*Kocher *et al.* (1994); Folch *et al.* (2010); Dehouck *et al.* (2009); Pucci *et al.* (2016)). More precisely, we used our recently developed solubility-dependent statistical potentials (Hou *et al.*, 2018) to discriminate between residue pair interactions that favor or disfavor protein solubility. In addition to these energetic features, we considered a series of other structure-based features and of commonly used sequence features. These were integrated into a predictor with the help of a random forest regression algorithm, so as to predict protein solubility with improved accuracy. Our predictor, called SOLart, is made freely available online at http://babylone.ulb.ac.be/SOLART/.

## 2. Materials AND Methods

### 2.1. Protein solubility definition

We used as a definition of solubility 𝒮 (in %) the ratio of the supernatant fraction obtained after centrifugation of the translation mixture over the total concentration of the overexpressed protein (Niwa *et al.*, 2009). It ranges from 0% to 130%. It is generally different from the physical solubility 𝒮_0_, measured in g/l and defined as the concentration of protein in a saturated solution that is in equilibrium with a solid phase.

𝒮_0_ is difficult to measure and strongly depends on the type of precipitant used to perform the experiment and on the environmental variables such as the temperature. This makes the construction of a large dataset of 𝒮_0_ values for training and testing bioinformatic models almost impossible. We thus chose to use the 𝒮 definition that can be measured and studied in large-scale investigations (Niwa *et al.*, 2009; Uemura *et al.*, 2018).

### 2.2. Protein datasets

To train and test SOLart, we considered two datasets of proteins that were expressed with the cell-free expression system called PURE (Shimizu *et al.*, 2005) and whose solubilities 𝒮 were measured. These are Esol_*Ecoli*_ and Esol_*Scerevisiae*_, which contain the solubilities of about 70% of the entire *E. coli* K-12 strain proteome (Niwa *et al.*, 2009), and of around 500 cytosolic proteins from *S. cerevisiae* (Uemura *et al.*, 2018), respectively.

We used the functional and structural annotation server EcoGene (Zhou and Rudd, 2013) and the UNIPROT server (Apweiler *et al.*, 2004) to map the gene accession ids of every entry in these datasets onto the corresponding structures from the Protein Data Bank (PDB) (Berman *et al.*, 2000). Only X-ray structures with maximum 2.5 Å resolution, which have a sequence identity of 100% and at least 90% coverage with the associated Esol sequences, were selected.

We also considered the remaining proteins from the Esol datasets, which have no experimental structure. We collected structural models for these entries from the SWISS-MODEL repository (Bienert *et al.*, 2016). Only the models constructed using a template with a good resolution X-ray structure (*≤* 2.5 Å) and at least 30% sequence identity and 50% coverage with the query sequence were kept.

We first focused on the set of X-ray structures from *E. coli*. We used the protein-culling server PISCES (Wang and Dunbrack Jr, 2003) to select proteins with pairwise sequence identity of 25% at most. This dataset, called 𝒟_*Ecoli*_, contains 406 well resolved protein structures with experimental solubility values and low pairwise sequence identity. It is used as the SOLart training set.

Out of the modeled structures from *E. coli*, we dropped those that have a sequence identity of more than 40% with a protein from 𝒟_*Ecoli*_, and filtered out sequences with more than 40% pairwise identity. We obtained in this way the ℳ_*Ecoli*_ dataset containing 679 protein models, which were used as a first test set.

We used the same procedure on the datasets of X-ray and modeled structures from *S. cerevisiae* proteins: we removed the entries that have more than 40% sequence identity with the training set 𝒟_*Ecoli*_, and finally filtered out proteins with more than 40% pairwise sequence identity. In this way, we obtained a third test set 𝒟_*Scerevisiae*_ composed of 70 X-ray structures and a fourth test set ℳ_*Scerevisiae*_ with 64 structures obtained via homology modeling.

The proteins that are contained in the four datasets are listed in Tables S1 of Supplementary Material, with some additional information.

Our datasets could be suspected to be biased and to contain only some types of conformations. To check that they do not suffer from this problem, we mapped all structures from our datasets onto the corresponding CATH categories (Dawson *et al.*, 2016). As shown in Table 1, the 406 X-ray structures of 𝒟_*Ecoli*_ belong to 344 homologous superfamilies, which cover all four classes, 59% of the architectures and 15% of the folds. The three test datasets 𝒟_*Scerevisiae*_, ℳ_*Ecoli*_ and ℳ_*Scerevisiae*_ do not cluster into the same superfamilies. This indicates that the protein sets are unbiased, and not specifically enriched in certain superfamilies but tend to cover the full fold universe.

**Table 1.** Mapping of the proteins of our datasets onto CATH categories (Dawson *et al.*, 2016). The numbers in parentheses correspond to the amount of proteins in the corresponding dataset. ‘C’ stands for Class, ‘A’ for Architecture, ‘T’ for Topology or fold, and ‘H’ for Homologous superfamily. ‘n’ is the number of C, A, T or H categories that have at least member in the dataset and ’%’ the fraction of these categories represented in the dataset.

| | $\mathcal{D}_{Ecoli}$<br>(406) | | $\mathcal{M}_{Ecoli}$<br>(679) | | $\mathcal{D}_{Scerevisiae}$<br>(70) | | $\mathcal{M}_{Scerevisiae}$<br>(64) | |
| --- | --- | --- | --- | --- | --- | --- | --- | --- |
|  | n | % | n | % | n | % | n | % |
| C | 4 | 100 | 4 | 100 | 3 | 75 | 3 | 75 |
| A | 24 | 59 | 26 | 63 | 15 | 37 | 15 | 37 |
| T | 214 | 15 | 244 | 18 | 55 | 4 | 54 | 4 |
| H | 344 | 6 | 458 | 7 | 70 | 1 | 64 | 1 |

## 3. Results

### 3.1. Features

We used a series of features to set up the SOLart solubility predictor, which are described below.

#### • Statistical potentials

We applied and extended the solubility-dependent statistical potentials recently introduced in (Hou *et al.*, 2018), which have proven to yield an objective and informative description of the interactions that modulate protein solubility properties. The idea was to divide the dataset 𝒟_*Ecoli*_ into two subsets of equal size, called 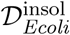 and 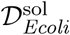, which contain aggregation-prone and soluble proteins, respectively, and to derive distance potentials from each of the the two subsets (see (Hou *et al.*, 2018) for details). In this way, we defined two distinct potentials referred to as ”insoluble” and ”soluble”.

Here we generalized this construction to other types of potentials involving various sequence and structure elements. In particular, for potentials based on one sequence element *s* and one structure element *c*, we have:

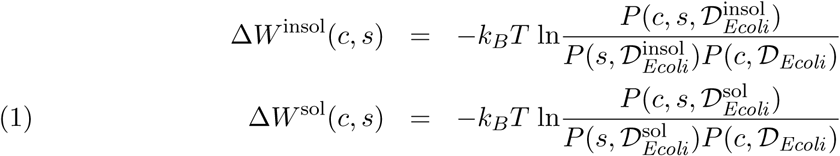

where *k*_*B*_ is the Boltzmann constant and *T* the absolute temperature. The sequence descriptor *s* is an amino acid type, and the structure descriptor *c* is either an inter-residue distance *d* computed between the average geometric centers of the heavy side chain atoms, a backbone torsion angle domain *t*, or a solvent accessibility *a* computed as the ratio between the solvent accessibility of a residue in a given structure and in an extended Gly-X-Gly tripeptide conformation (see *e.g.* (Kocher *et al.*, 1994; Rooman *et al.*, 1991; Pucci *et al.*, 2014)). *P* (*s, c*, 𝒟) is the probability of joint occurrence of the sequence and structure elements *s* and *c* in the dataset 𝒟, and similarly for the probability functions *P* (*s*, 𝒟) and *P* (*c*, 𝒟). These probabilities were estimated in terms of the number of occurrences of the sequence-structure associations in 𝒟.

We constructed eleven solubility-dependent statistical potentials from different combinations of *s* and *c* elements, listed in Table 2. We named the potentials according to the type and number of sequence and structure descriptors. For example, ”sa” represents the potential in which one amino acid type and one solvent accessibility are specified, whereas ”sds” describes the potential in which two amino acid types and their interresidue distance are given.

**Table 2.** List of all the features tested for SOLart. Those used in the final version are marked by a 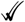; those for which a subset is used are marked by a ✓.

| Features | Description | SOLart |
| --- | --- | --- |
| **Statistical potentials** |  |  |
| sd: $\Delta\Delta G_{sd}$ | 1 amino acid, 1 distance | ✓✓ |
| sds: $\Delta\Delta G_{sds}$ | 2 amino acids, 1 distance | ✓✓ |
| sa: $\Delta\Delta G_{sa}$ | 1 amino acid, 1 solvent accessibility | ✓✓ |
| saa: $\Delta\Delta G_{saa}$ | 1 amino acid, 2 solvent accessibilities | ✓✓ |
| ssa: $\Delta\Delta G_{ssa}$ | 2 amino acids, 1 solvent accessibility | ✓✓ |
| st: $\Delta\Delta G_{st}$ | 1 amino acid, 1 torsion angle domain | ✓✓ |
| stt: $\Delta\Delta G_{stt}$ | 1 amino acid, 2 torsion angle domains | ✓✓ |
| sst: $\Delta\Delta G_{sst}$ | 2 amino acids, 1 torsion angle domain | ✓✓ |
| sad: $\Delta\Delta G_{sad}$ | 1 amino acid, 1 distance and 1 solvent accessibility | ✓✓ |
| std: $\Delta\Delta G_{std}$ | 1 amino acid, 1 distance and 1 torsion angle domain | ✓✓ |
| sta: $\Delta\Delta G_{sta}$ | 1 amino acid, 1 distance and 1 solvent accessibility | ✓✓ |
| **Protein size and solvent accessible surface area** |  |  |
| $\Lambda$ | protein length | ✓✓ |
| SAcc | protein solvent accessibility | ✓✓ |
| SAcc/$\Lambda$ | protein solvent accessibility divided by length | ✓✓ |
| **Secondary structure content** |  |  |
| $\beta\_b$ | fraction of buried $\beta$ residues | ✓✓ |
| $\beta\_m$ | fraction of moderately buried $\beta$ residues | ✓✓ |
| $\beta\_e$ | fraction of exposed $\beta$ residues | |
| $\alpha\_b$ | fraction of buried $\alpha$ residues | |
| $\alpha\_m$ | fraction of moderately buried $\alpha$ residues | ✓✓ |
| $\alpha\_e$ | fraction of exposed $\alpha$ residues | ✓✓ |
| $\gamma\_b$ | fraction of buried coil residues | |
| $\gamma\_m$ | fraction of moderately buried coil residues | |
| $\gamma\_e$ | fraction of exposed coil residues | |
| **Amino acid composition** |  |  |
| $C_i$ ($i=1..20$) | fraction of each of the 20 amino acid types | ✓ |
| K+R | fraction of positively charged residues |  |
| K-R | fraction of K minus fraction of R | ✓✓ |
| D+E | fraction of negatively charged residues | ✓✓ |
| D-E | fraction of D minus fraction of E |  |
| K+R+D+E | fraction of charged residues | ✓✓ |
| K+R-D-E | fraction of positively minus negatively charged residues | ✓✓ |
| F+W+Y | fraction of aromatic residues | ✓✓ |
| $\_b, m, e$ | *idem* with distinction between buried, moderately buried and exposed residues | ✓ |

We used these different potentials to compute folding free energy contributions of target proteins. As an example, the folding free energies computed from the soluble and insoluble ssd distance potentials were defined as:

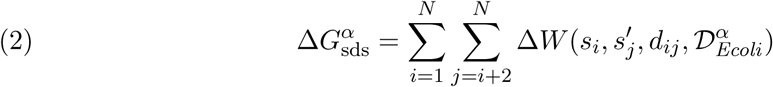

where *α* is equal to ”sol” or ”insol”, *s*_*i*_ and *s*^*′*^_*j*_ are two residue types at positions *i* and *j* along the sequence, *d*_*ij*_ is their spatial distance, and *N* is the number of amino acids of the target protein. We then computed the folding free energy difference:

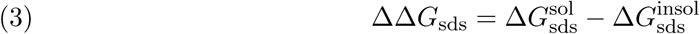

Using analogous relations, we computed the folding free energy Δ*G*^*α*^ and the folding free energy difference ΔΔ*G* for each potential listed in Table 2.

#### • Protein size and accessible surface area

We considered three global characteristics of the proteins, which are the protein length (Λ), its solvent accessible surface area (SAcc) estimated with an in-house program (Dalkas *et al.*, 2014), and its solvent accessible surface area divided by the protein length (SAcc/Λ); in the latter case we used the length of the sequence whose structure has been determined. Note that the former feature is sequence-based, and that the latter two require the knowledge of the 3D structure.

#### • Secondary structure content

Another series of structure-based features were added, which are the fraction of protein residues that are in *α*-helical, *β*-strand or coil (called here *γ*) conformation. We distinguished between the *α, β* and *γ* residues that are buried in the protein core (solvent accessibility *≤* 20%), moderately buried (between 20 and 50%), and solvent exposed (*≥* 50%). Our in-house program (Dalkas *et al.*, 2014) was used to assign the secondary structure and solvent accessibility.

#### • Amino acid composition

We integrated 20 purely sequence-based features, corresponding to the fraction of each of the 20 amino acid present in a protein. We also considered the fraction of amino acid groups, *i.e.* positively charged residues (K+R), negatively charged residues (D+E), charged residues (K+R+D+E) aromatic residues (F+W+Y), as well as the difference between the fractions of K and R (K-R), D and E (D-E), and K+R and D+E (K+R-D-E). We combined these features with the solvent accessibility and defined three categories per amino acid or amino acid group, according to whether the residue is exposed, moderately buried or buried. This yielded 81 additional structure-based features.

### 3.2. Feature selection

The next step consisted in selecting, out of the above-defined 28 purely sequence-based features and 103 structure-based features, the subset of features that are the most informative for protein solubility. We used for that purpose the 𝒟_*Ecoli*_ training set, which contains 406 non-redundant high-resolution X-ray structures of *E. coli* proteins with low pairwise sequence identity and experimentally measured solubility (see Methods). The feature selection was performed using the Boruta algorithm (Kursa *et al.*, 2010) implemented in the Caret package of R (Kuhn *et al.*, 2008), a wrapper built around the random forest classification algorithm (Liaw *et al.*, 2002), which compares the importance of the real features with those of random (shadow) features using statistical testing. The results are obtained as an average over several runs (here 1,000) of random forest.

We filtered out the features whose average importance measured by the Boruta algorithm is lower than 1. This led us to keep a total of 52 features, which are shown in in Fig. 1 and Supplementary Information Fig. S1. Among these, 37 require the knowledge of the structure.

**Figure 1.**
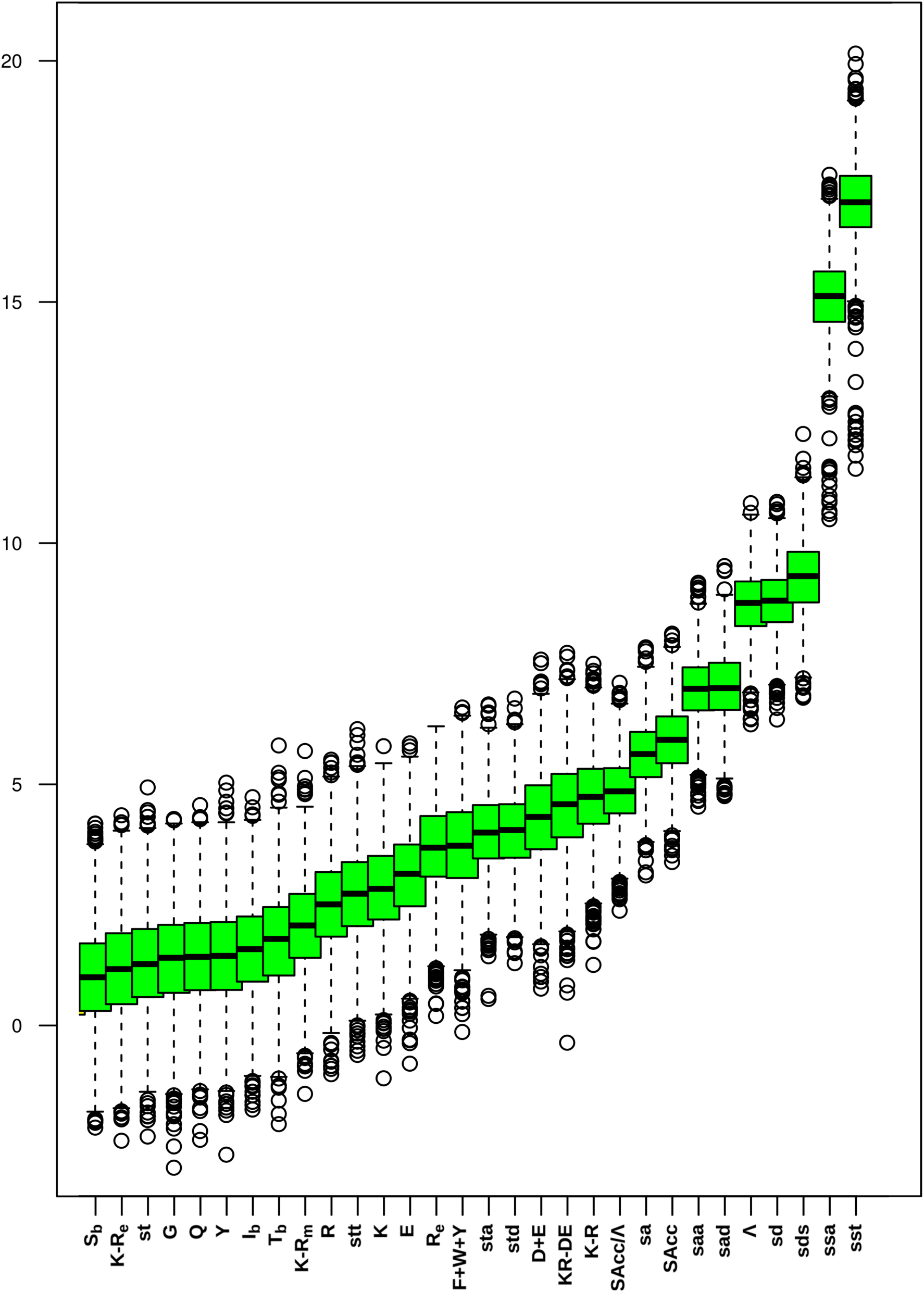
The top 30 most important features identified by feature selection, from left to right. The names in lower-case letters indicate folding free energy differences, *e*.g sst means ΔΔ*G*_sst_.

Strikingly, the four top-ranked features are folding free energy differences ΔΔ*G* computed from our solubility-dependent potentials: the backbone torsion angle potential sst, the solvent accessibility potential ssa and the two distance potentials sd and sds (see Table 2). The next most important feature is the protein length Λ, followed by the solvent accessibility and fractions of some amino acid types. The features based on the secondary structure do not appear among the 30 top features, but some appear in the list of 52 selected features.

### 3.3. Setting up SOLart

The 52 selected features were combined to set up the SOLart predictor of the solubility of target proteins on the basis of their 3D structures. We used for that purpose 𝒟_*Ecoli*_ as training set, and the random forest regression algorithm implemented in the R package (Liaw *et al.*, 2002) to construct the model. This algorithm is a tree-based system composed of multiple regression trees; the number of trees is here set to 500. The training process starts with a randomly selected subset of the original dataset from which a regression tree is constructed by the iterative partitioning of the data space into smaller subsets. At each node of the tree, randomly sampled features are used; the number of features depends on a global parameter ”mtry” taken here between 1 and 3. The optimal mtry value is obtained through a grid search procedure as the one that yields the highest correlation coefficient in the training dataset. The regression for a target protein is obtained by averaging the predictions over all trees.

### 3.4. Performance of SOLart

SOLart’s performances were evaluated by three replicates of a 10-fold cross validation procedure on the 𝒟_*Ecoli*_ training set. The replicates were performed with different random divisions into folds, and the performances were computed as averages over the replicates. Our computational model reaches a good linear correlation coefficient *r* = 0.67 between the SOLart solubility predictions and the experimental values, and a root mean square error RMSE= 25% (Table 3).

**Table 3.** **SOLart performances** in cross validation on the learning set 𝒟_*Ecoli*_, and on three independent test sets: 𝒟_*Scerevisiae*_ containing X-ray structures and ℳ_*Ecoli*_ and ℳ_*Scerevisiae*_ containing modeled structures. The values in parentheses correspond to the performance with 10% outliers removed.

| | $\mathcal{D}_{Ecoli}$ | $\mathcal{M}_{Ecoli}$ | $\mathcal{D}_{Scerevisiae}$ | $\mathcal{M}_{Scerevisiae}$ |
| --- | --- | --- | --- | --- |
| $r$ | 0.67 | 0.51 (0.66) | 0.70 (0.78) | 0.65 (0.71) |
| RMSE | 25% | 28% (23%) | 23% (19%) | 24% (20%) |

We also tested SOLart on an independent test set that contains *S. cerevisiae* proteins with a well resolved X-ray structure, grouped in the 𝒟_*Scerevisiae*_ set (see Methods). The performance of SOLart on this set is evaluated by a linear correlation coefficient *r* = 0.70 and an RMSE = 23%. When 10% outliers are removed, the score increases up to *r* = 0.78 and RMSE = 19% (Table 3). The scores on this independent set are thus even better than those obtained in cross validation on the training set 𝒟_*Ecoli*_.

To further analyze this result, we estimated the importance of each feature in the SOLart prediction using the varImp permutation scheme-based function (Kuhn *et al.*, 2008). It proceeds by randomly permuting each feature in turn in order to break its association with the response, and then using it together with the remaining unpermuted features for prediction. The decrease of the prediction accuracy is a measure of the importance of the permuted feature. This measure estimates the weight of each individual feature in the predictor, whereas the feature selection algorithm applied in section 3.2 measures the feature relevance independently of the prediction model. They thus yield slightly different rankings.

The 20 most important features of our prediction model are shown in Fig. 3. Interestingly, almost all the features that correspond to folding free energy differences (ΔΔ*G*) are in this list (9 out of 11), and the 6 top features are the ΔΔ*G*s computed from the potentials ssa, sst, sd, sds, saa, and sa (Table 2). The two best ones, almost *ex æquo*, are ΔΔ*G*_ssa_ and ΔΔ*G*_sst_, which also ranked first in the feature selection (Fig. 1). They are computed from the propensities of amino acid pairs to be associated with a certain solvent accessibility range *a* or a certain backbone torsion angle domain *t* of a residue. These propensities differ between soluble and aggregation-prone proteins, and it is this difference which is measured through the ΔΔ*G* features. The next best ranked features are ΔΔ*G*_sd_ and ΔΔ*G*_sds_, computed from the propensities of residue pairs to be separated by a certain spatial distance, followed by two other accessibility potentials ΔΔ*G*_saa_ and ΔΔ*G*_sa_.

These folding free energy features require the protein structure as input. In fact, more than half of the top 20 features are structure-based. This confirms the relevance of the structural information in the determination of the protein solubility properties. The first sequence-based feature ranks seventh. It is the sequence length Λ: in general, the smaller the sequence, the most soluble the protein (Kramer *et al.*, 2012). The two related features, *i.e.* the solvent accessible surface area SAcc divided or not by the length, are also among the top 20 features.

**Figure 2.**
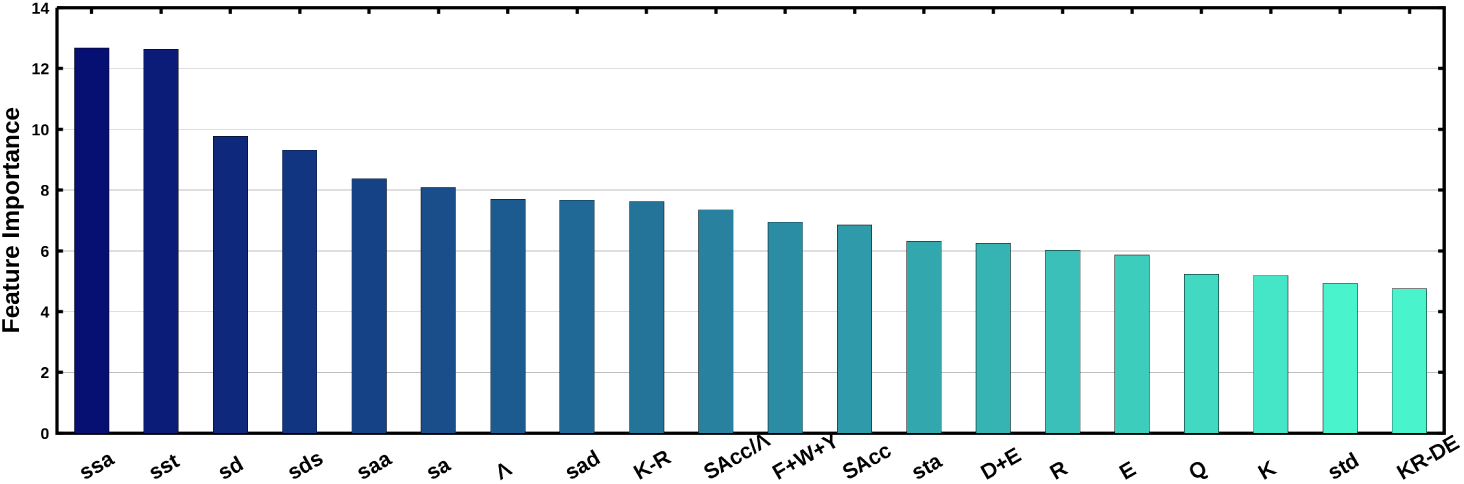
The top 20 most important features of SOLart, from right to left. The names in lower-case letters indicate folding free energy differences, *e*.g sst means ΔΔ*G*_ssa_.

**Figure 3.**
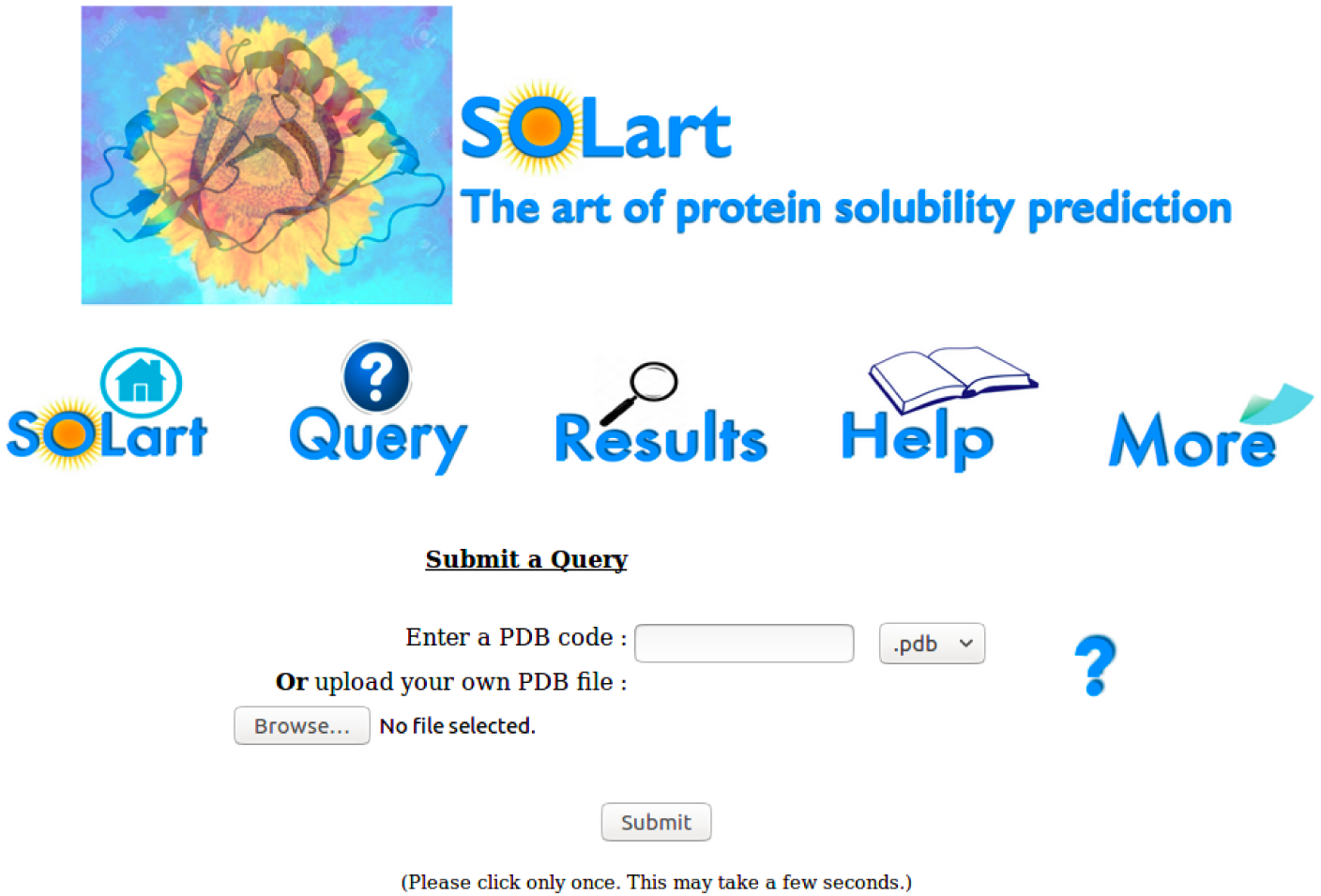
The webserver interface of SOLart.

The remaining top 20 features are sequence-based: the difference between Lys and Arg composition (K-R) which is positively correlated with solubility (Warwicker *et al.*, 2013; Hou *et al.*, 2018), the percentage of aromatic residues (F+Y+W) which favor aggregation (Niwa *et al.*, 2009; Hou *et al.*, 2018), and the total fraction of negatively charged residue (D+E) that have also been shown to promote solubility (Niwa *et al.*, 2009; Hou *et al.*, 2018). The next features are the composition in R and Q, which disfavors solubility, the composition in E and K, which instead promotes solubility, and the difference between the fraction of positively and negatively charged residues (K+R-D-E), which augments insolubilty.

Note that all these sequence-based features are also employed by the solubility predictors available in the literature. However, in addition to these commonly used features, we exploit a series of structure-based features among which the most important ones are obtained from the newly developed solubility-dependent statistical potentials. These capture the solubility properties in a more accurate way and represent the key instrument of our approach.

### 3.5. Performance on modeled protein structures

SOLart has been shown to be accurate when the 3D structure of the target protein is known. To enlarge its applicability, we tested it on low-resolution structures obtained via homology modeling. We first applied it to the ℳ_*Ecoli*_ dataset containing 679 proteins from *E. coli* (see Methods). We obtained a correlation of *r* = 0.51 and a RMSE of 28%, which is relatively good but lower than the performance on 𝒟_*Ecoli*_ (Table 3). This drop is expected since we have to take into account the possible inaccuracies in the modeled structures that have to be added to the error of our computational method. After removing 10% outliers, the performance increases to *r* = 0.66 and RMSE= 23%, and reaches thus the same performance as on good-resolution structures.

As a last test set, we used ℳ_*Scerevisiae*_ that contains *S. cerevisiae* proteins with modeled structures. The performance of SOLart on this set is given by *r* = 0.65 and RMSE = 24%, and increases up to *r* = 0.71 and RMSE = 20% without 10% outliers. The scores are thus much higher on this test set than on the *E. coli* test set, which suggests that some structural protein models or experimental solubility values might be less accurate on the the *E. coli* set than on the *S. cerevisiae* set.

Note that these tests are quite strict, since there is a low sequence similarity (¡40%) between these test sets and the training set. We thus conclude that SOLart can reliably be used to predict solubility not only for high-resolution experimental structures but also for modeled or other low-resolution structures.

### 3.6. Comparison with other solubility prediction methods

The performance of SOLart was compared with that of other solubility prediction methods on the combination of 𝒟_*Scerevisiae*_ and ℳ_*Scerevisiae*_ sets, that group X-ray and modeled structures from *S. cerevisiae* proteins, as these are independent test sets that are not included in the training sets of any of the predictors. More precisely, we tested the methods Protein-SOL (Heb-ditch *et al.*, 2017), ccSOL (Agostini *et al.*, 2014), CamSol (Sormanni *et al.*, 2015), PROSO (Smialowski *et al.*, 2006), PROSO II (Smialowski *et al.*, 2012) and SOLpro (Magnan *et al.*, 2009), by submitting to their respective webservers all the proteins from our test datasets. Note that all these methods are sequence-based.

The linear correlation coefficient *r* between the solubility predictions and the experimental values for all these predictors are given in Table 4. Our method clearly outperforms the competitors (*r* = 0.68 against *r* = 0.56 for the second best method). This demonstrates the importance of using structural information.

**Table 4.** Comparison of the performance of different predictors on the combination of the 𝒟_*Scerevisiae*_ and ℳ_*Scerevisiae*_ test sets, on the basis of the Pearson correlation coefficient between predicted and experimental solubility values. SOLpro and PROSO first predict the proteins as soluble or insoluble and then give a probability score; we thus calculated the correlation by considering the solubility to be -1 for proteins predicted as insoluble and +1 for proteins predicted as soluble.

| Predictor | $r$ |
| --- | --- |
| SOLart | 0.68 |
| Protein-Sol | 0.56 |
| ccSOL | 0.56 |
| CamSol | 0.39 |
| PROSO II | 0.13 |
| SOLpro | 0.18 |
| PROSO | 0.20 |

### 3.7. Webserver

We provided a freely available webserver interface for our prediction method, which targets non-expert users (http://babylone.ulb.ac.be/SOLART/index.php) (Fig. 4). The input consists of the 3D structure of the target protein in PDB format. It can be uploaded directly by the user or imported from the PDB (Berman *et al.*, 2000) by typing its 4-letter code. The webserver then provides a brief summary of some of the protein’s characteristics and allows the user to choose one of the protein chains. The computation starts after the query submission. All the structure-based free energy, secondary structure and solvent accessibility features are first computed and then integrated with the other, sequence-based, features.

**Figure 4.**
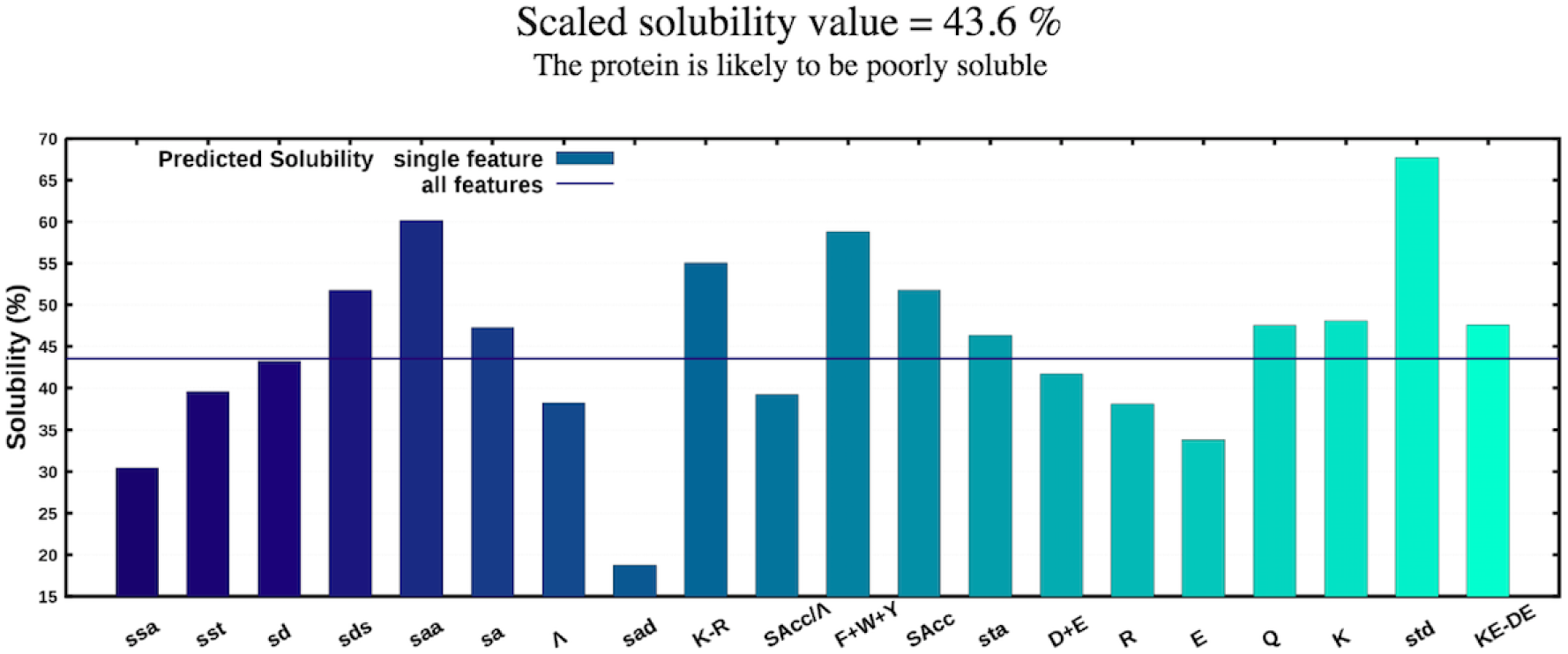
Predicted solubility of an example protein (PDB code 2qia, Uniprot code P0A722) with all features used in SOLart (horizontal line) or with each single feature only (histogram bars).

In the output page, reached by following the link provided, the value of the predicted scaled solubility 𝒮 is given. If the score is close to zero, the target protein is predicted as aggregation-prone and, when it is close to 130, as soluble. Moreover, to have an indication of the contribution of each single feature to the solubility prediction of the target protein, we also show a figure (Fig. 4) with the solubility predicted from each feature taken individually and with all SOLart features together. The prediction with each single feature is computed from a random forest model trained on the experimental solubility values of the 𝒟_*Ecoli*_ set. This figure can be used as a source of inspiration to suggest the characteristics to modify in view of modulating solubility.

Due to its simplicity of use, we expect that this webserver will be of interest for researchers in academia and industry who are interested in modulating protein solubility without needing any prior bioinformatic knowledge.

## 4. Discussion

We introduced SOLart as on of the first structure-based solubility prediction method, which is able to predict quickly and accurately the protein solubility of a protein from its experimental or modeled 3D structure.

SOLart employs a series of features, among which the sequence-based features that are commonly used for solubility prediction and some classical structure-based features such as secondary structure composition and solvent accessibility. In addition, it takes advantage of the potentiality of solubility-dependent statistical potentials to discriminate the residue interactions that favor aggregation or solubility. Besides the distance potentials that have previously been analyzed (Hou *et al.*, 2018), ten new solubility-dependent potentials were introduced here, which describe the local propensities of residues to adopt certain backbone torsion angle domains or to have certain solvent accessibility values in soluble or aggregation-prone proteins. Note that the feature importance analyses show that the torsion, solvent accessibility and distance potentials are the most important features in the random forest regression prediction. The folding free energy differences computed with these potentials are better correlated with solubility than other protein properties analyzed in the literature such as protein length, isoelectric point and aliphatic index.

The performances of SOLart are high and robust: the linear correlation on both the training dataset (in cross validation) and on three independent test sets almost reaches 0.7 on good-resolution X-ray structures and slightly lower on modeled structures. It is important to underline that SOLart can be used with modeled structures, as it largely expands the domain of applicability of our tool. Furthermore, it performs similarly in the training and testing datasets, which indicates its robustness and absence of bias towards the training set. Finally, SOLart outperforms the state-of-the-art solubility predictors on an independent dataset containing *S. Cerevisiae* proteins, with an increase of more thant 20% in the correlation coefficient between the predicted and the experimental values of the solubility. This provides a strong demonstration of SOLart’s accuracy and usefulness.

Another advantage of SOLart is its fastness: it is able to predict the solubility of a medium-size protein in less than one minute. This quality make this tool a perfect instrument to investigate protein solubilty properties on a large scale.

Even though SOLart performances are good, there is still a lot of work needed to unravel the various effects and to understand the biophysical mechanisms underlying solubility and aggregation. One direction is to design better energy functions that describe more efficiently these phenomena by enlarging the protein datasets with experimental solubility values or modifying their original formulation. For example, the definition of the reference state that is adequate for solubility properties is still an open problem. It has been argued that interactions between unfolded conformations could lead to insoluble aggregates and, indeed, inclusion bodies forming in heterologous expression in *E. coli* have been shown to involve folded, unfolded, misfolded and partially folded proteins (Martínez-Alonso *et al.*, 2009; Singh *et al.*, 2015; Baneyx and Mujacic, 2004; Singh and Panda, 2005; Vallejo and Rinas, 2004), which makes it challenging to disentangle the characteristics contributing to its formation.

Note also that the definition of the solubility (𝒮) used in this paper differs from the physical definition of solubility (𝒮_0_), measured in g/l, defined as the concentration of a protein in a saturated solution that is in equilibrium with a solid phase. To get insights into the relation between these two solubility definitions, they should systematically be compared. This is currently impossible as no large datasets of 𝒮_0_ values are available due to the difficulties in its experimental measurements.

A final perspective concerns industrial biotechnological applications, in which water is replaced by other polar solvents or even by non-polar solvents. Understanding how the protein solubility changes according to the type of solvent and being able to accurately predict this change is a major target for computational tools. On the same footing, it would also be important to understand and predict the influence of buffer salts and ionic strength on the solubility properties of proteins.

In summary, SOLart is a new and efficient method to predict protein solubility. Thanks to its user-friendly interface, both expert and non-expert users can use its predictions to analyze and improve the solubility properties of targeted proteins involved in biotechnological processes, where solubility is frequently a major bottleneck.

